# Lack of ITS sequence homogenization in congeneric plant species with different ploidy levels

**DOI:** 10.1101/2022.05.29.493735

**Authors:** Carolina Osuna-Mascaró, Rafael Rubio de Casas, Modesto Berbel, José M. Gómez, Francisco Perfectti

## Abstract

The internal transcribed spacers (ITS) exhibit concerted evolution by the fast homogenization of these sequences at the intragenomic level. However, the rate and extension of this process are unclear and might be conditioned by the number and divergence of the different ITS copies. In some cases, such as hybrid species and polyploids, ITS sequence homogenization appears incomplete, resulting in multiple haplotypes within the same organism. Here, we studied the dynamics of concerted evolution in 85 individuals of seven plant species of the genus *Erysimum* (Brassicaceae) with multiple ploidy levels. We estimated the rate of concerted evolution and the degree of sequence homogenization separately for ITS1 and ITS2 and whether these varied with ploidy. Our results showed incomplete sequence homogenization, especially for polyploid samples, indicating a lack of concerted evolution in these taxa. Homogenization was usually higher in ITS2 than in ITS1, suggesting that concerted evolution operates more efficiently on the former. Furthermore, the hybrid origin of several species appears to contribute to the maintenance of high haplotype diversity, regardless of the level of ploidy. These findings indicate that sequence homogenization of ITS is a dynamic and complex process that might result in varying intra- and inter-genomic diversity levels.

## Introduction

Concerted evolution is an evolutionary process by which sequences from the same gene family show higher sequence similarity to each other than to orthologous genes in related species [1, 2]. Hence, genes evolved in a concerted manner present low polymorphism in their sequences, i.e., the sequences are homogenized. Concerted evolution is particularly notable in multicopy nuclear genes, where homogenization is mainly achieved by unequal crossing over and gene conversion [3, 4]. One of the best characterized multicopy gene families is the 45S nuclear ribosomal DNA (nrDNA). It appears arranged as tandem repeated units with hundreds to thousands of copies in one or several loci per genome. These units are composed of the 18S rDNA, internal transcribed spacer 1 (ITS1), 5.8 S rDNA, internal transcribed spacer 2 (ITS2), and 26S rDNA, separated by longer non-transcribed intergenic spacers [5]. Among all these units, the internal transcribed spacers (ITS1 and ITS2) are the best-characterized nrDNA sequences [6] partly because ITS sequences show characteristics advantageous for phylogenetic studies, such as biparental inheritance, short length, and high evolution rate [4, 7, 8].

ITS sequences usually present fast concerted evolution with low levels of intra-genomic sequence variation and very few polymorphic positions [9, 10]. However, in some animals (e.g., [11]) and especially in plants [12, 13, 14, 15, 16, 17], sequence homogenization remains incomplete across ITS sequences, resulting in relatively high intra-genomic polymorphism. This ITS diversity is often linked to hybridization events [8, 18, 19, 20, 21]. Different ITS sequences may meet after hybridization and become homogenized after a time, but this homogenization may not be consistent among descendant lineages [22]. As concerted evolution tends to homogenize sequences rapidly [8], evidence of non-concerted evolution is mostly expected in recently-formed hybrid species, where both parental ITS sequences may still be present. This phenomenon should be particularly conspicuous in recent allopolyploid species, where the occurrence of different ITS sequences located in distinct chromosomes tends to delay this homogenization [23].

*Erysimum L*. (Brassicaceae) comprises more than 200 species [24], mainly from Eurasia, with some species inhabiting North America and North Africa [25, 26]. The Baetic Mountains (SE Iberia) constitute one of the most important glacial refugia in Europe and a hotspot for this group, with ∼10 *Erysimum* species occurring in this small area [27, 28]. Previous studies have suggested that several of these taxa could have a hybrid origin [29, 30, 31]. Ploidy levels vary among and, in some cases, within species [27, 32], suggesting that a detailed understating of hybridization and allopolyploidization is necessary to shed light on the evolution of this group. However, the effects of hybridization and polyploidization on the genomes of these species are far from being fully understood.

In this study, we explore the homogenization dynamics of ITS, taking into account the interacting effects of concerted evolution, hybridization, and polyploidization. For this purpose, we analyzed polymorphisms at the species, population, and individual levels in ITS1 and ITS2 for seven *Erysimum* species. We sequenced both markers by NGS to recover all the ITS copies present in the different genomes [10, 33]. With these sequences, we then proceeded to quantify the degree of sequence homogenization in both ITS1 and ITS2 within individuals, populations, and species; and the concerted evolution levels in polyploid *Erysimum* spp. Our results showed heterogeneous levels of ITS sequence homogenization with reduced concerted evolution, especially for polyploids. Any insight into ITS evolution in plants needs to consider the concomitant effects of hybridization and polyploidization on the rates of concerted evolution.

## Materials and methods

### Taxon sampling

We collected fresh leaves from polyploid and diploid *Erysimum* species (see Table 1 for details on species ploidy levels). In particular, we sampled five individuals from three different populations of *Erysimum baeticum, E. bastetanum, E. mediohispanicum, E. nevadense*, and *E. popovii*, and five individuals from one population of *E. lagasca*e and five from the microendemic *E. fitzii*. A total of 85 samples (= individuals) were dried and preserved in silica gel until DNA extraction. Table 1 shows the code, location, and ploidy levels of all samples.

**Table 1.**
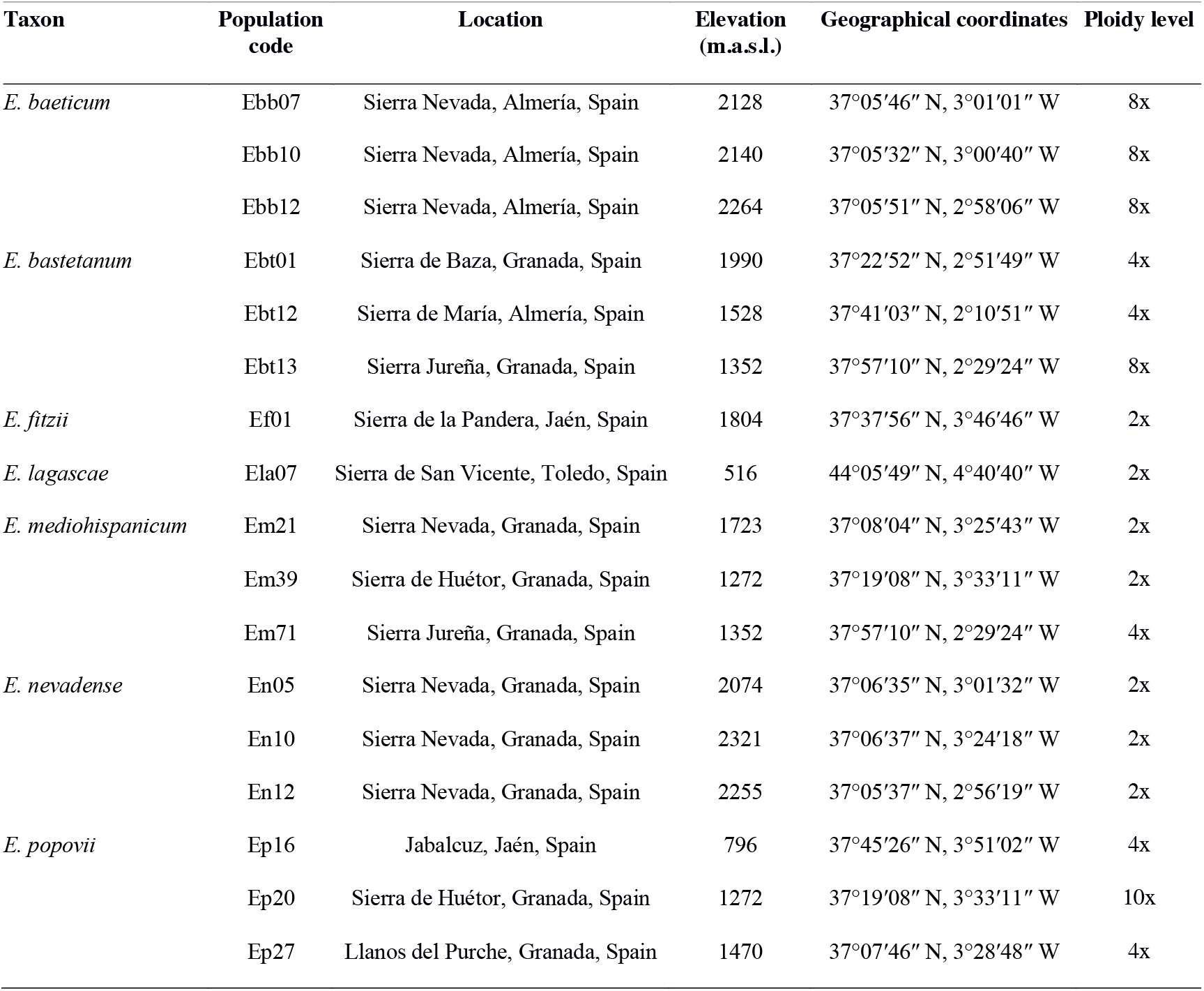
Taxonomic assignment, population code, location, elevation, and ploidy level for the *Erysimum* spp. populations sampled.

### DNA extraction

We used at least 60 mg of dry plant material from each sample. We disrupted the tissues in liquid N2 using a mortar and pestle. Then, total genomic DNA was isolated using the GenElute Plant Genomic DNA Miniprep kit (Sigma-Aldrich, St. Louis, MO) following the manufacturer’s protocol. The quantity and quality of the obtained DNA were checked using a NanoDrop 2000 spectrophotometer (Thermo Fisher Scientific, Wilmington, DE, United States), and the integrity of the extracted DNA was checked on agarose gel electrophoresis.

### ITS1 and ITS2 amplification

We independently amplified ITS1 and ITS2 in each sample. The ITS PCR reactions were performed in 25 μl with the following composition: 5 μL 5× buffer containing MgCl2 at 1.5 mM (New England Biolabs), 0.1 mM each dNTP, 0.2 µM each primer, and 0.02 U Taq high fidelity DNA-polymerase (Q5 High-Fidelity DNA Polymerase, New England Biolabs). We used a set of long primers developed to have a 5’ flanking sequence complementary to the Nextera XT DNA index to facilitate adapter ligation during library construction:

>ITS1-Flabel

TCG TCG GCA GCG TCA GAT GT GTA TAA GAG ACA GTC CGT AGG TGA ACC TGC GG

>ITS2-Rlabel

GTC TCG TGG GCT CGG AGA TGT GTA TAA GAG ACA GGC TGC GTT CTT CAT CGA TGC

>ITS3-Flabel

TCG TCG GCA GCG TCA GAT GTG TAT AAG AGA CAG GCA TCG ATG AAG AAC GCA GC

>ITS4-Rlabel

GTC TCG TGG GCT CGG AGA TGT GTA TAA GAG ACA GTC CTC CGC TTA TTG ATA TGC

Reactions included 30 cycles with the following conditions: 94 °C 15 s, 60 °C 30 s, and 72 °C 30 s. Amplified fragments were purified using spin columns (GenElute TM PCR Clean-Up Kit, Sigma-Aldrich) and were checked on agarose gel electrophoresis. Finally, we quantified the starting DNA concentration using the Infinite M200 PRO NanoQuant spectrophotometer (TECAN, Männedorf, Switzerland).

### Library construction

We constructed two libraries, one for ITS1 amplicons and one for ITS2 amplicons. The libraries were prepared using the Nextera XT DNA Sample Preparation Kit. In brief, the DNA was tagged by adding a unique adapter label combination to the 3’ and 5’ ends of the DNA sequence. Then, the DNA was amplified via a nine-cycle PCR. The total volume reaction was 25 μl with the following composition: 5 μL 10× buffer at 1.0 mM (New England BioLabs), 0.1 mM each dNTP, 0.2 µM each Nextera primer, 0.02 U Taq high fidelity DNA-polymerase (Q5, NEB), and 5× Q5 High GC Enhancer (NEB). PCR thermocycling conditions were 98 °C during 5 s, 55 °C for 10 s, and 72 °C for 10 s. After that, we purified both libraries using the GenElute PCR Clean-Up Kit (Sigma) to remove short library fragments. Finally, we generated equal volumes of the libraries to prepare equimolar libraries for sequencing, and the final concentration of each library was quantified using the Infinite M200 PRO NanoQuant spectrophotometer (TECAN, Männedorf, Switzerland).

### Library sequencing

ITS1 and ITS2 library sequencing were carried out by Novogene Bioinformatics Technology Co., Ltd, with an Illumina MiSeq platform (Illumina, USA) using a paired-end 150 bp sequence read run. The ITS libraries of *E. mediohispanicum* were sequenced twice due to an unexpected low sequencing output (we constructed new libraries as explained above). This sequencing was done using the Illumina Miseq platform and paired-end chemistry in the Center for Scientific Instrumentation (CIC) of the University of Granada, Spain.

## Data analysis

FASTQ files were demultiplexed, and read quality was checked in FastQC v0.11.5 [34]. Then, we did a trimming procedure using first cutadapt v1.15 [35] to trim the adapters, followed by a quality trimming using Sickle v1.33 [36]. Forward and reverse reads were paired in Geneious R.11 [37]. Using the function “Set pair read” with default parameters for Illumina paired-end read technology. Then the paired reads were merged using BBMerge v37.64 [38] with a “Low Merge rate” to decrease false positives. Then, to reduce redundancy and noise caused by sequencing errors and tag switching events, we did a cluster analysis using CD-HIT v4.6.8 [39]. We clustered the sequences from each sample using an identity threshold of 0.99 (i.e., we merged the sequences with similarity ≥ 99%) and discarded the clusters that included < 5 % of the total reads [40]. This step reduced the contribution of sequencing errors to the reported sequence diversity.

We aligned the sequences from each sample using MAFFT v7.450 [41] with default parameters, generating one alignment per species and marker. We trimmed the alignments using trimAl v1.2 [42], removing poorly aligned regions with the “gappyout” method. We estimated population genetic parameters at intra-species, intra-population, and intra-individual levels using the R package PEGAS v0.1 [43]. We used the “nuc.div” function to calculate nucleotide diversity (π), estimated as the average number of nucleotide differences per site between two sequences [44, 45]. Moreover, we estimated the haplotype diversity (Hd), with the “hap.div” function, as the probability of differentiation between two randomly chosen haplotypes. We then used the “haplotype” function to calculate the total number of haplotypes and the haplotype frequency distribution for each species, population, and individual. We checked for normality using Shapiro-Wilk’s method and then compared the nucleotide and haplotype diversity and the number of haplotypes among polyploid and diploid species and among ITS1 and ITS2 using the Mann-Whitney-Wilcoxon test. All statistical analyses were done in R v 4.1.0 using the package stats v3.6.1 [46].

We investigated potential correlations among ploidy levels and haplotype and nucleotide diversity for ITS1 and ITS2 samples. Also, as these species were described as frequently hybridizing, we studied if there were shared haplotypes among different populations of the same species and among different species. To explore that, we estimated the total number of ITS1 and ITS2 haplotypes and their frequencies.

We analyzed the genetic structure of ITS1 and ITS2 by performing a hierarchical analysis of molecular variance (AMOVA; [47]). We used the “amova” function from the R package PEGAS v0.1 [43] to explore the genetic variation explained by populations (i.e., at the population level), among individuals within populations (i.e., at the individual level), and within individuals (i.e., at the intra-genome level). Moreover, we analyzed the amount of genetic variation in ITS1 and ITS2 explained by interspecific differences by partitioning the variance into three levels: among species, among populations within species, and within populations (i.e., among individuals).

## Results

From the initial 85 individuals, we obtained good-quality sequences for a total of 84 ITS1 and 81 ITS2 samples, with 10,156 ± 1,233 sequences per individual for ITS1 and 49,428 ± 7,678 sequences for ITS2 (Table S1).

Polyploid species (*E. baeticum, E. bastetanum, E. popovii*) tended to have higher nucleotide diversity than diploid species (Figure 1) for both ITS1 (Wilcoxon test = 655, p-value: 0.04; mean π ± SE; polyploid: 0.012 ± 0.007, diploid: 0.004 ± 0.006) and ITS2 (Wilcoxon test = 663, p-value: 0.03; polyploids: 0.003 ± 0.004, diploids: 0.002 ± 0.003). In addition, the polyploid population of *E. mediohispanicum* (Em71, 4x) showed higher nucleotide diversity than the two diploid populations of this species, marginally significant for ITS1 (Wilcoxon test = 10, p-value: 0.05; Em71: mean π = 0.011 ± 0.006; Em39: mean π = 0.006 ± 0.007 for ITS1; Em21: mean π = 0.0004 ± 0.001; Figure S1) and more pronounced for ITS2 (Wilcoxon test = 8.5, p-value: 0.04; Em71: mean π = 0.003 ± 0.002; Em39: mean π = 0.0003 ± 0.001 for ITS2; Em21: mean π = 0.001 ± 0.002 for ITS2; Figure S1). Furthermore, the correlation between ploidy level and nucleotide diversity was highly significant for ITS1 (Spearman’s rho: 0.48, p-value: 2.10×10^−6^; Figure S2) and marginally significant for ITS2 (Spearman’s rho: 0.20, p-value: 0.06). The difference in the degree of association of ITS1 and ITS2 polymorphisms with ploidy levels might be a consequence of overall diversity. ITS2 samples presented significantly lower nucleotide diversities than ITS1 ones (Wilcoxon test = 5165.5, p-value: 3.33×10^−7^). Nucleotide diversity values for ITS1 and ITS2 at the three levels of analysis (species, population, individual) are shown in Tables S2-S8.

**Figure 1.**
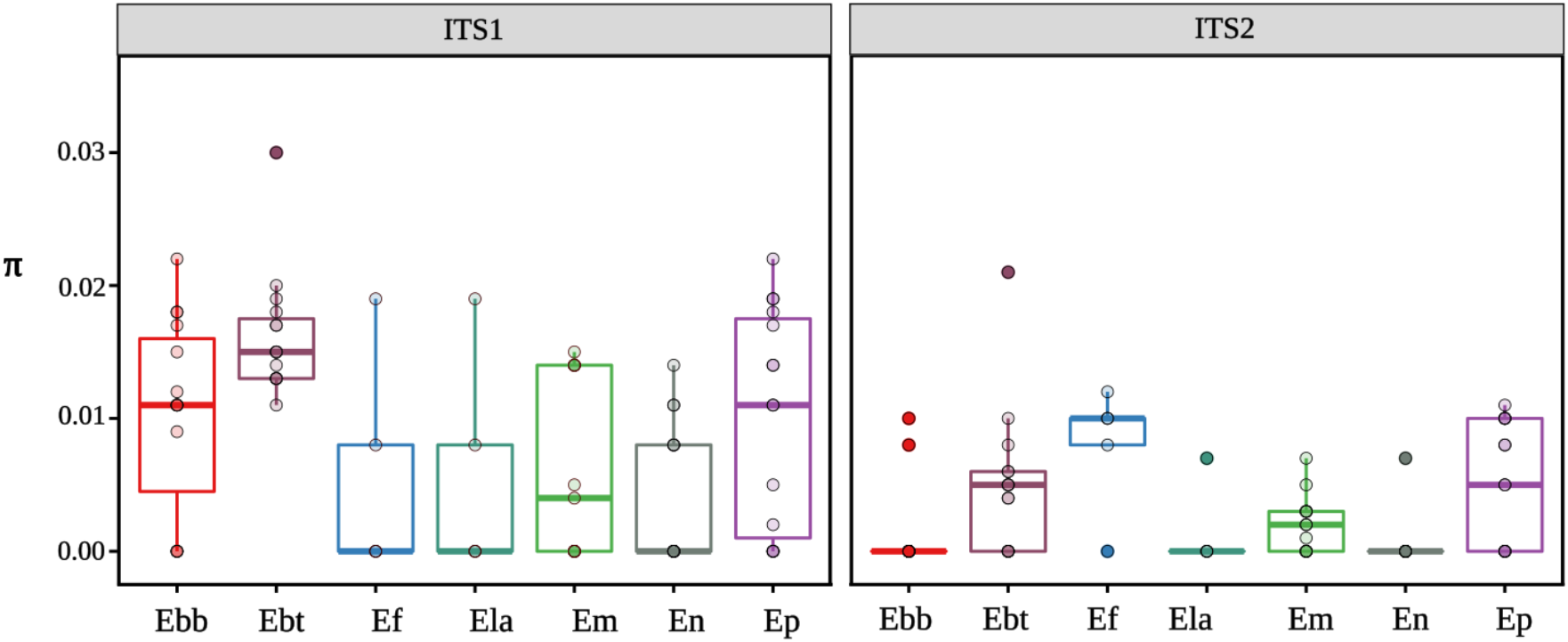
Boxplot depicting the nucleotide diversity (π) for ITS1 and ITS2 samples. Nucleotide polymorphism was estimated for each *Erysimum* individual as the average number of nucleotide differences per site between two sequences (Nei and Li, 1979). *E. baeticum* (Ebb), *E. bastetanum* (Ebt), *E. popovii* (Ep), and one population of *E. mediohispanicum* (Em) are polyploids. *E. nevadense* (En), *E. fitzii* (Ef), two populations of *E. mediohispanicum* (Em), and *E. lagascae* (Ela) are diploids.

**Figure 2.**
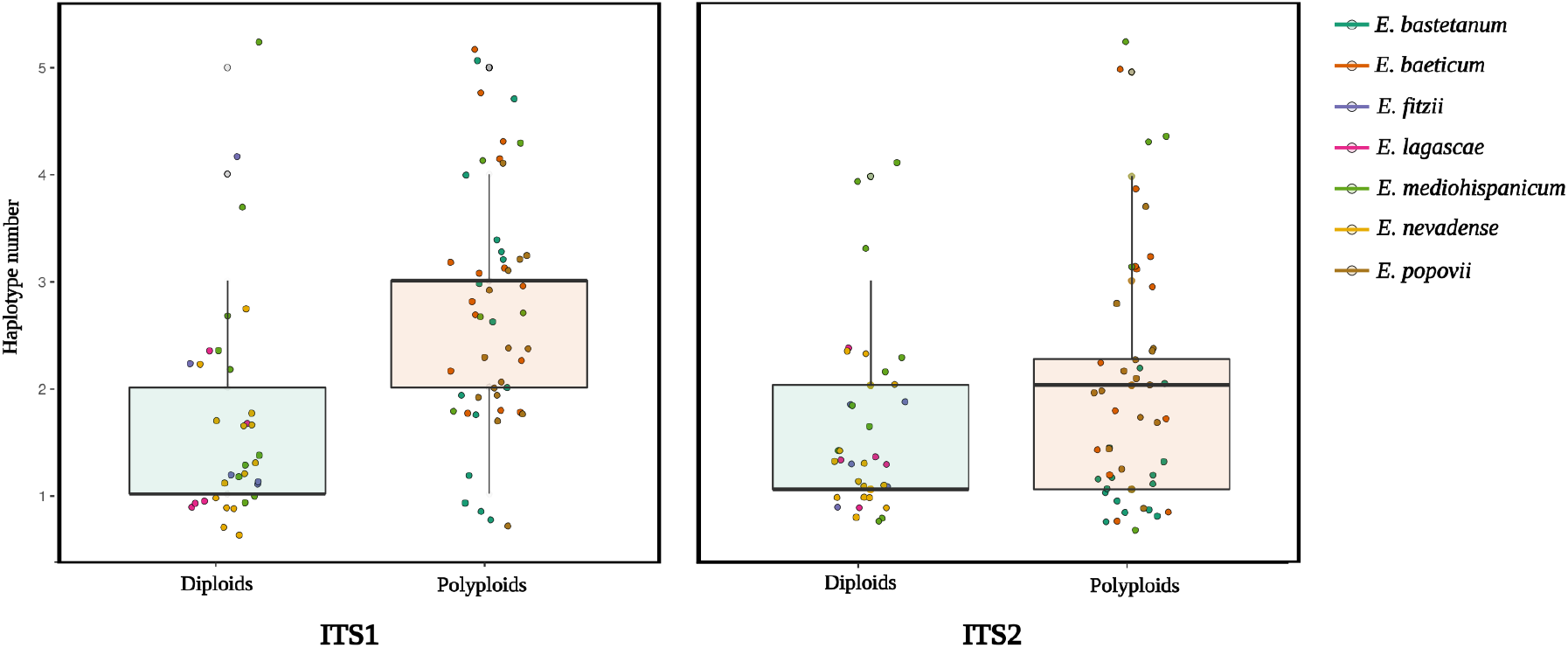
Boxplot depicting the number of ITS1 (left) and ITS2 (right) haplotypes per sample for diploid and polyploid species.

Haplotype diversity showed a similar pattern to that of the nucleotide diversity, with higher haplotype diversity for polyploid species than diploid species, for ITS1 (Wilcoxon test = 343, p-value: 2.16×10^−6^; mean Hd = 0.89 ± 0.38 for polyploid; mean Hd = 0.50 ± 0.49 for diploid) and marginally significant for ITS2 (Wilcoxon test = 632.5, p-value = 0.059; mean Hd = 0.39 ± 0.49 for polyploid; mean Hd = 0.28 ± 0.45 for diploid). Moreover, the degree of association between haplotype diversity and ploidy level seemed to differ between ITSs, being highly significant for ITS1 (Spearman’s rho: 0.43, p-value: 2.96×10^−5^) but only marginally significant for ITS2 (Spearman’s rho: 0.18, p-value: 0.09). The values of haplotype diversity for both ITS and three levels are shown in Tables S2-S8. ITS2 presented lower haplotype diversity than ITS1 in terms of haplotype numbers (Wilcoxon test = 4458, p-value 0.002; Table 2). ITS2 diversity was reduced to a single haplotype (i.e., no polymorphism was detected) in 49 individuals (Table S2-S8). Conversely, only 30 individuals showed no nucleotide diversity in ITS1 (Table S2-S8).

**Table 2.** Average haplotype diversity (Hp) per species, estimated for ITS1 and ITS2 samples. Maximum and minimum values (in parentheses) refer to individual samples.

| Species | ITS 1 | ITS2 |
| --- | --- | --- |
| *E. baeticum* | 0.983<br>(1 - 0.963) | 0.897<br>(1 - 0.933) |
| *E. bastetanum* | 0.983<br>(1 - 0.969) | 0.893<br>(1 - 0.666) |
| *E. fitzii* | 0.944<br>(1 - 0) | 0.872<br>(1 - 0) |
| *E. lagascae* | 0.936<br>(1 - 0) | 0.733<br>(1 - 0) |
| *E. mediohispanicum* | 0.969<br>(1 - 0.400) | 0.805<br>(1 - 0.866) |
| *E. nevadense* | 0.938<br>(1 - 0.785) | 0.941<br>(1 - 0.833) |
| *E. popovii* | 0.984<br>(1 - 0.888) | 0.943<br>(1 - 0.866) |

Several ITS1 haplotypes were shared across species, particularly among some populations of *E. bastetanum, E. fitzii, E. mediohispanicum*, and *E. nevadense*. Specifically, we found that the three populations of *E. bastetanum* studied in this article shared haplotypes with two *E. mediohispanicum* populations (Em39, Em71) and with the three populations of *E. nevadense*. In addition, *E. bastetanum* populations and one population of *E. nevadense* (En05) shared haplotypes with the *E. fitzii* population included in the analyses. Conversely, no ITS2 haplotypes were found to be shared across different species (Tables S10, S11, S13, and S14).

The hierarchical AMOVA showed that interspecific differences were a significant source of variation for both ITS (Table 3). The species-level explained 52.63 % and 73.50 % of the variance for ITS1 (p-value < 0.001, Φ = 0.48) and ITS2 (p-value < 0.001, Φ = 0.70) respectively, implying ample genetic divergence among species. Conversely, differences among populations were not significant and absorbed a relatively low amount of molecular variance (< 9% for both ITS1 & ITS2; Table 3). When the genetic structure was separately analyzed for each species, we found more complex results. Most of the variance (44.96 % – 100 % for ITS1; 29.12 % – 100 % for ITS2) resided within-individuals (see Table 4). Differences among populations varied from 0 % to 48.07 % for ITS and from 0 % to 70.87 % for ITS2. Moreover, the differences were only significant in *E. mediohispanicum, E. nevadense*, and *E. popovii* for ITS1 and *E. bastetanum* for ITS2 (Table 4).

**Table 3.** Hierarchical AMOVA results for ITS1 and ITS2 regions.

| Sequence | Source of variation | df | Variance (sigma<sup>2</sup>) | % Variance | Φ statistics | p-value |
| --- | --- | --- | --- | --- | --- | --- |
| ITS1 | Species | 6 | 1.96 x 10<sup>-5</sup> | 52.63 | 0.48 | < 0.01 |
|  | Populations within species | 10 | 2.23 x 10<sup>-6</sup> | 6.00 | 0.55 | 0.58 |
|  | Within populations | 197 | 1.54 x 10<sup>-5</sup> | 41.36 | - | - |
| ITS2 | Species | 6 | 1.10 x 10<sup>-4</sup> | 73.50 | 0.7 | < 0.01 |
|  | Populations within species | 10 | 1.32 x 10<sup>-5</sup> | 8.84 | 0.8 | 0.99 |
|  | Within populations | 128 | 2.64 | 17.64 | - | - |

**Table 4.** ITS1 and ITS2 hierarchical AMOVA results for *E. baeticum, E. bastetanum, E. fitzii, E.lagascae, E. mediohispanicum, E. nevadense*, and *E. popovii*.

| Species | Source of variation | ITS1 |  |  |  |  | ITS2 |  |  |  |  |
| --- | --- | --- | --- | --- | --- | --- | --- | --- | --- | --- | --- |
|  |  | df | Variance (sigma<sup>2</sup>) | Variance (%) | Φ | p | df | Variance (sigma<sup>2</sup>) | Variance (%) | Φ | p |
| *E. baeticum* | Populations | 2 | 1.48 x 10<sup>-5</sup> | 8.93 | 0.11 | 0.09 | 2 | 1.38 x 10<sup>-7</sup> | 0.3 | 0.01 | 0.46 |
|  | Individuals within populations | 12 | 0 | 0 | 0 | 0.99 | 12 | 0 | 0 | 0 | 0.99 |
|  | Within individuals | 24 | 1.51 x 10<sup>-4</sup> | 91.06 | - | - | 2 | 4.46 x 10<sup>-5</sup> | 99.69 | - | - |
| *E. bastetanum* | Populations | 2 | 0 | 0 | 0 | 0.96 | 2 | 8.05 x 10<sup>-5</sup> | 70.87 | 0.75 | <0.01 |
|  | Individuals within populations | 12 | 0 | 0 | 0 | 0.61 | 10 | 0 | 0 | 0.69 | 0.99 |
|  | Within individuals | 31 | 2.33 x 10<sup>-4</sup> | 100 | - | - | 29 | 3.30 x 10<sup>-5</sup> | 29.12 | - | - |
| *E. fitzii* | Individuals within populations | 4 | 8.37 x 10<sup>-5</sup> | 55.03 | 0 | 0.14 | 3 | 0 | 0 | 0 | 0.9 |
|  | Within individuals | 4 | 6.48 x 10<sup>-5</sup> | 44.96 | - | - | 5 | 6.08 x 10<sup>-5</sup> | 100 | - | - |
| *E. lagascae* | Individuals within populations | 4 | 6.50 x 10<sup>-8</sup> | 0.07 | 0 | 0.55 | 4 | 0 | 0 | 0 | 0.94 |
|  | Within individuals | 2 | 8.48 x 10<sup>-5</sup> | 99.92 | - | - | 4 | 0.06 | 100 | - | - |
| *E. mediohispanicum* | Populations | 2 | 4.58 x 10<sup>-5</sup> | 48.07 | 0.5 | <0.01 | 2 | 0 | 0 | 0 | 0.97 |
|  | Individuals within populations | 12 | 0 | 0 | 0.45 | 0.69 | 12 | 0 | 0 | 0 | 0.28 |
|  | Within individuals | 42 | 4.95 x 10<sup>-5</sup> | 51.92 | - | - | 10 | 5.08 x 10<sup>-5</sup> | 100 | - | - |
| *E. nevadense* | Populations | 2 | 1.67 x 10<sup>-5</sup> | 18.29 | 0.25 | <0.01 | 2 | 0 | 0 | 0 | 0.99 |
|  | Individuals within populations | 11 | 0 | 0 | 0 | 0.8 | 11 | 0 | 0 | 0 | 0.88 |
|  | Within individuals | 7 | 7.47 x 10<sup>-5</sup> | 81.7 | - | - | 3 | 5.12 x 10<sup>-5</sup> | 100 | - | - |
| *E. popovii* | Populations | 2 | 3.13 x 10<sup>-5</sup> | 19.01 | 0.2 | 0.02 | 2 | 1.15 x 10<sup>-3</sup> | 7.73 | 0.09 | 0.56 |
|  | Individuals within populations | 12 | 0 | 0 | 0.13 | 0.72 | 12 | 0 | 0 | 0 | 0.41 |
|  | Within individuals | 20 | 1.33 x 10<sup>-4</sup> | 80.98 | - | - | 12 | 0.01 | 92.26 | - | - |

## Discussion

We observed incomplete sequence homogenization for the 45S rDNA regions in the *Erysimum* species studied here. Our analyses were based on stringent trimming to avoid false polymorphisms due to sequencing errors. However, despite being so restrictive, we found high nucleotide and haplotype diversities overall, especially for ITS1, and a significant genetic structure that may inform the evolutionary history of these species.

Polyploid *Erysimum* species presented lower ITS homogenization levels than diploid species. Specifically, polyploid species presented higher nucleotide and haplotype diversity and a higher number of haplotypes, congruent with the hypothesis that polyploids harbor greater genetic diversity even within gene families [48]. The lack of concerted evolution in polyploid species has been previously described in several plant species in which an absence of sequence homogenization could be related to a recent allopolyploid origin [33, 49, 50, 51, 52]. Moreover, some studies have suggested that the number of rDNA loci, usually located in different chromosomes, is expected to be higher in polyploids, hindering sequence homogenization [53, 54]. The number of rDNA loci and their chromosomic locations in these *Erysimum* species is unknown. In the genome of the diploid *E. cheiranthoides* [55], the rDNA appears in eight locations in chromosomes 3, 6, 7, and 8, which may be related to the number of rDNA loci for the diploid *Erysimum* species studied here. In any case, a relatively higher number of rDNA loci is expected for polyploid *Erysimum* species. Although the number of rDNA loci may coincide with the sum of those of its parents in young allopolyploids, it could be more variable in older polyploids, where some loci are usually lost [56, 57, 58, 59].

We also detected limited sequence homogenization in diploid species, particularly in ITS1. The high molecular variance within diploid genomes (Tables 4 and 5) could be the result of past hybridization events, which might result in the coexistence of multiple ITS families within individual genomes, particularly if hybrids are young [33, 60]. This result is congruent with previous studies, in which the genomes of the diploid *Erysimum* species studied here were found to exhibit signatures of recent hybridization and introgression [32]. Moreover, *Erysimum* phylogenies based on ITS sequences [61, 62, 63] showed a variable degree of phylogenetic incongruence compatible with hybridization. Here, the influence of hybridization on ITS diversity is further supported by the significant molecular variance among populations detected in some species (i.e., *E. mediohispanicum, E. nevadense*, for ITS1), showing a non-consistent homogenization pattern in the population level. Thus, these results suggest a different history of hybridization for each population, in concordance with previous studies [31, 32].

Our results indicated that sequence homogenization was heterogeneous across the 45S rDNA regions within a general scenario of high diversity. The degree of polymorphism exhibited by ITS1 was much higher than that of ITS2, suggesting that concerted evolution is operating more efficiently on the latter. This result agrees with previous studies that have shown that ITS1 is, on average, more variable than ITS2, which has been described as a very conserved marker [64, 65, 66, 67, 68, 69]. This variation between the two spacers might help to analyze evolutionary patterns at different scales. While ITS1 variation might throw light on divergence at the population- or individual-level, our AMOVA results (Table 3) suggest that ITS2 could be useful for species-level characterization, at least in *Erysimum* spp.

Because of their sensitivity to hybridization, ITS markers have been previously used to identify the parental contributors of hybrid taxa [16, 49, 60, 70, 71]. Our study found shared haplotypes among diploid and polyploid species (specifically among *E. bastetanum* – a polyploid – and the diploid species *E. fitzii, E. mediohispanicum*, and *E. nevadense*), which could be the result of incomplete lineage sorting or the effect of recent hybridization events. However, we have not found decisive evidence of whether these diploid species could be considered parental species of the polyploid taxon. Moreover, our results indicate hybridization across taxonomic levels (i.e., from individuals to species) since they are more congruent with multiple backcrossings across populations and taxa than with a single, “original” allopolyploidization event. Reticulated evolution seems to be the norm in this genus [32, 55, 72, 73, 74, 75]. Thus, for the species analyzed in this study, Osuna-Mascaro et al. (2022) [32] have found genomic evidence of rampant introgression between species, including both lilac- and yellow-flowered species. Future studies identifying the alleles co-located on the same chromosome through phased haplotypes [76] or using PacBio single-molecule sequencing and the PURC method (Pipeline for Untangling Reticulate Complexes; [77]) could be used to identify parental species of the different hybrid taxa and trace back the evolutionary patterns of these *Erysimum* species.

Despite their evident versatility as molecular evolution markers, the analysis of ITS sequences needs to be undertaken to realize that concerted evolution might often be insufficient to ensure sequence homogenization [78]. Both ITS and, especially, ITS2 have for a long time been used as phylogenetic and barcoding markers in plants [7, 65, 79, 80, 81]. However, many studies have pointed out that evolutionary inferences based on these markers might lead to misleading or erroneous conclusions in species where sequence homogenization is lacking due to hybridization or other genome rearrangement events [8, 10]. In this study, our results indicate that allopolyploidization and hybridization have severely impaired ITS sequence homogenization in *Erysimum*, implying that ITS-based phylogenies of this genus should be considered with prudence. Given that these causes of genomic rearrangement are widespread and prevalent among flowering plants [10], caution is advised when using ITS for phylogenetic studies without prior knowledge of haplotype distribution, even for diploid species. Hence, intragenomic variation for ITS sequences could be used as an indication of possible recent hybridization.

## Acknowledgments

The authors thank the Tatina López Pérez and the Evoflor group for helping us during several phases of the study. We also thank the Sierra Nevada National Park headquarters for providing access to sampling in the National Park. This research is supported by grants from FEDER/Junta de Andalucía-Consejería de Economía y Conocimiento A-RNM-505-UGR18 and P18-FR-3641. This research was also initially funded by the Spanish Ministry of Science and Innovation (CGL2013-47558-P), including EU FEDER funds. COM was supported by the Ministry of Economy and Competitiveness (BES-2014-069022). This is a contribution to the Research Unit Modeling Nature, funded by the Consejería de Economía, Conocimiento, Empresas y Universidad, and European Regional Development Fund (ERDF), reference QUALIFICA 00011.

## Author contributions

COM, RR, and FP conceived and designed the study. COM and MB carried out the laboratory procedures and field sampling, with the help of FP and JMG. COM analyzed the data with the help of FP. COM wrote the first draft. The final version of the M.S. was redacted with the contribution of all the authors.

## Data availability

The raw data from this project were submitted to NCBI Sequence Read Archive (SRA) and can be found by the BioSample ID: SUB11440702, BioProject PRJNA835881, and the following accession numbers: *E. baeticum* (Ebb07: SAMN28120146, Ebb10: SAMN28120147, Ebb12: SAMN28120148); *E. bastetanum* (Ebt01: SAMN28120149, Ebt12: SAMN28120150, Ebt13: SAMN28120151); *E. fitzii:* Ef01 (SAMN28120152); *E. lagascae* (Ela07: SAMN28120153); *E. mediohispanicum* (Em21: SAMN28120154, Em39: SAMN28120155, Em71: SAMN28120156); *E. nevadense* (En05: SAMN28120157, En10: SAMN28120158, En12: SAMN28120159); *E. popovii* (Ep16: SAMN28120160, Ep20: SAMN28120161, Ep27: SAMN28120162).

## Competing interest

The authors declare no competing interests.

## Notes

### Competing Interest Statement

The authors have declared no competing interest.

