## Supplementary Material for "Lack of ITS sequence homogenization in congeneric plant species with different ploidy levels"

**Figure S1**. Boxplot depicting the nucleotide diversities (π) of ITS1 and ITS2 for *Erysimum mediohispanicum*.

**Figure S2.** Correlation between ploidy level and nucleotide diversity for ITS1.

**Table S1.** The number of sequences after quality trimming and after cd-hit clustering for all the samples.

**Table S2.** Nucleotide and haplotype diversity for *E. baeticum* ITS1 and ITS2, at the three-level analyzed.

**Table S3.** Nucleotide and haplotype diversity for *E. bastetanum* ITS1 and ITS2, at the three-level analyzed.

**Table S4.** Nucleotide and haplotype diversity for *E. fitzii* ITS1 and ITS2, at the two-level analyzed.

**Table S5.** Nucleotide and haplotype diversity for *E. lagascae* ITS1 and ITS2, at the three-level analyzed.

**Table S6.** Nucleotide and haplotype diversity for *E. mediohispanicum* ITS1 and ITS2, at the three-level analyzed.

**Table S7.** Nucleotide and haplotype diversity for *E. nevadense* ITS1 and ITS2, at the three-level analyzed.

**Table S8.** Nucleotide and haplotype diversity for *E. popovi* ITS1 and ITS2, at the three-level analyzed.

**Table S9.** Number of total haplotypes, frequency of each haplotype (based on the total of sequences after cd-hit analysis), number of haplotypes shared among different populations from the same species, and number of haplotypes shared among *E. baeticum* and other *Erysimum* species studied here.

**Table S10.** Number of total haplotypes, frequency of each haplotype (based on the total of sequences after cd-hit analysis), number of haplotypes shared among different populations from the same species, and number of haplotypes shared among *E. bastetanum* and other *Erysimum* species studied here.

**Table S11.** Number of total haplotypes, frequency of each haplotype (based on the total of sequences after cd-hit analysis), and number of haplotypes shared among *E. fitzii* and other *Erysimum* species studied here.

**Table S12.** Number of total haplotypes, frequency of each haplotype (based on the total of sequences after cd-hit analysis), and number of haplotypes shared among *E. lagascae* and other *Erysimum* species studied here.

**Table S13.** Number of total haplotypes, frequency of each haplotype (based on the total of sequences after cd-hit analysis), number of haplotypes shared among different populations from the same species, and number of haplotypes shared among *E. mediohispanicum* and other *Erysimum* species studied here.

**Table S14.** Number of total haplotypes, frequency of each haplotype (based on the total of sequences after cd-hit analysis), number of haplotypes shared among different populations from the same species, and number of haplotypes shared among *E. nevadense* and other *Erysimum* species studied here.

**Table S15.** Number of total haplotypes, frequency of each haplotype (based on the total of sequences after cd-hit analysis), number of haplotypes shared among different populations from the same species, and number of haplotypes shared among *E. popovii* and other *Erysimum* species studied here.

**Figure S1.** Boxplot depicting the nucleotide diversities (π) of ITS1 and ITS2 for *Erysimum mediohispanicum*. Populations Em21 and Em39 were diploids, and Em71 was polyploid (4×).

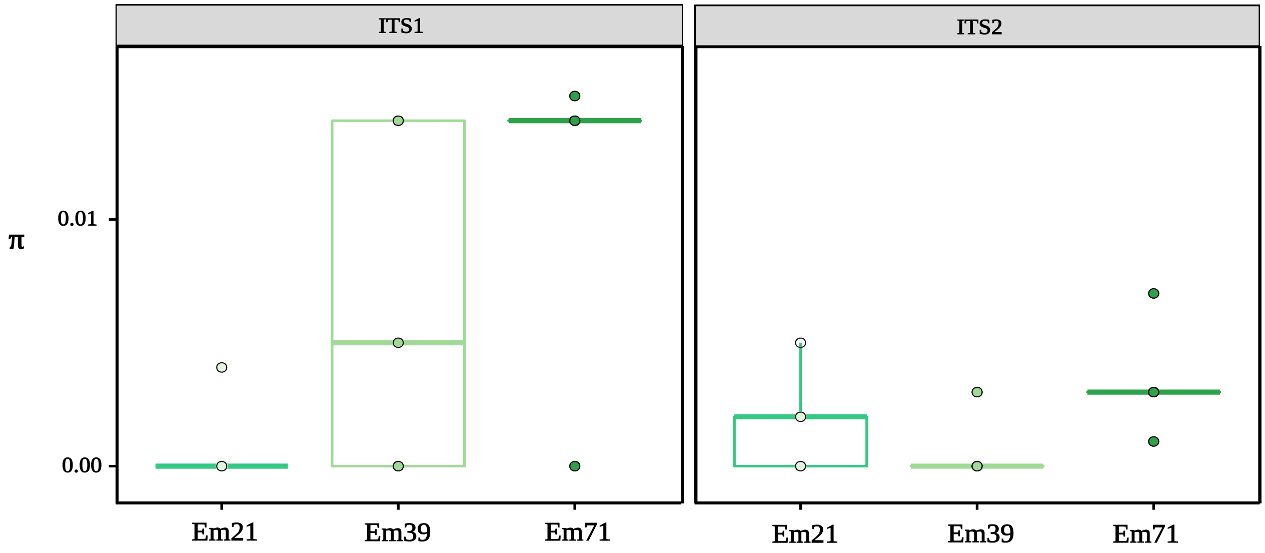

**Figure S2.** Correlation between ploidy level and nucleotide diversity for ITS1 samples (Spearman's rho: 0.48, p-value: 2.10 x $10^{-6}$). The ploidy level for the samples was: 2× (diploid), 4× (tetraploid), 8× (octoploid), and 10× (decaploid).

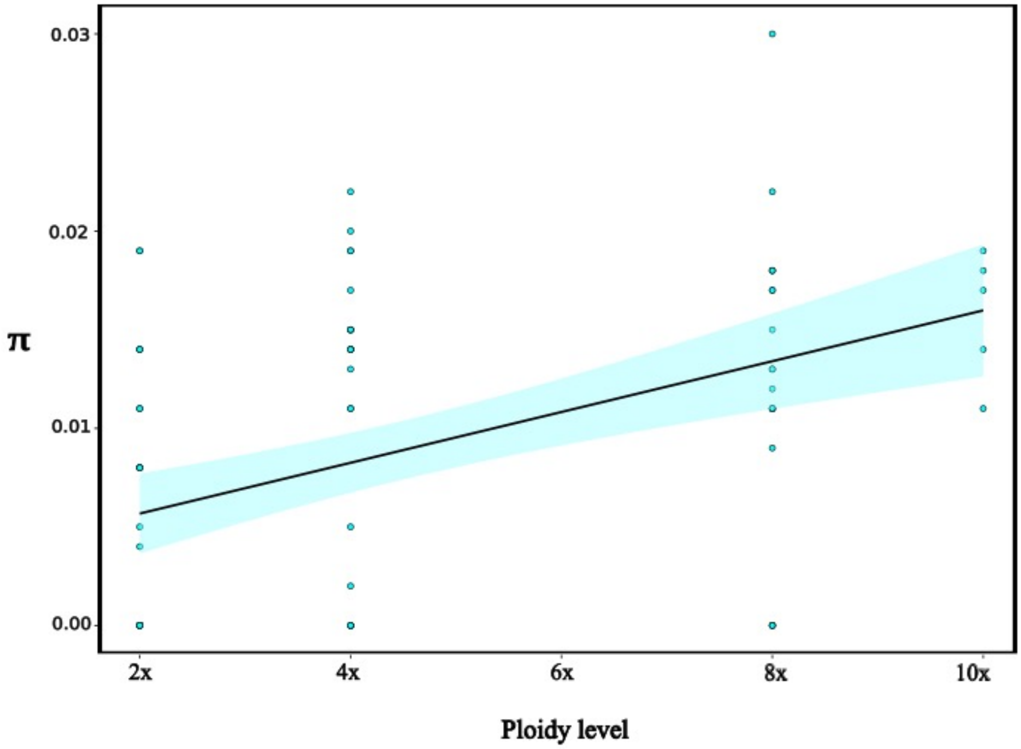

**Table S1.** Number of sequences after quality trimming and after cd-hit clustering for all the samples.

| Taxon | Sample | Number of sequences after quality trimming | | Number of sequences after clustering | |
|---|---|---|---|---|---|
| | | ITS1 | ITS2 | ITS1 | ITS2 |
| ***E. baeticum*** | Ebb07-1 | 9,323 | 194,745 | 7,122 | 191,686 |
| | Ebb07-2 | 7,358 | 25,941 | 6,539 | 25,311 |
| | Ebb07-3 | 5,852 | 45,680 | 5,274 | 18,641 |
| | Ebb07-4 | 5,198 | 334,388 | 3,731 | 132,263 |
| | Ebb07-5 | 6,092 | 403,632 | 5,063 | 128,485 |
| | Ebb10-1 | 7,090 | 608,256 | 5,289 | 247,990 |
| | Ebb10-2 | 9,843 | 181,762 | 9,171 | 65,007 |
| | Ebb10-3 | 6,710 | 181,726 | 6,155 | 109,565 |
| | Ebb10-4 | 9,468 | 352,662 | 8,618 | 152,423 |
| | Ebb10-5 | 10,847 | 381,902 | 9,190 | 160,503 |
| | Ebb12-1 | 9,820 | 452,562 | 8,586 | 172,469 |
| | Ebb12-2 | 6,487 | 600,370 | 5,328 | 262,497 |
| | Ebb12-3 | 6,706 | 441,938 | 5,727 | 167,879 |
| | Ebb12-4 | 6,013 | 493,274 | 5,403 | 201,674 |
| | Ebb12-5 | 8,379 | 71,674 | 6,784 | 29,745 |
| ***E. bastetanum*** | Ebt01-1 | 6,673 | 147,372 | 5,469 | 140,547 |
| | Ebt01-2 | 4,309 | 19,851 | 3,581 | 18,981 |
| | Ebt01-3 | 6,389 | 136,348 | 5,869 | 133,498 |
| | Ebt01-4 | 10,349 | 10,913 | 9,876 | 10,627 |
| | Ebt01-5 | 2,407 | 25,871 | 1,916 | 25,497 |
| | Ebt12-1 | 9,387 | 74,828 | 8,459 | 73,287 |
| | Ebt12-2 | 3,417 | 77,159 | 2,575 | 70,854 |
| | Ebt12-3 | 6,613 | 31,839 | 4,897 | 30,580 |
| | Ebt12-4 | 7,133 | 0 | 5,651 | 0 |
| | Ebt12-5 | 4,648 | 0 | 4,306 | 0 |
| | Ebt13-1 | 5,007 | 37 | 4,460 | 52 |
| | Ebt13-2 | 6,116 | 127 | 5,061 | 95 |
| | Ebt13-3 | 6,075 | 518 | 5,443 | 356 |
| | Ebt13-4 | 4,898 | 180 | 4,279 | 90 |
| | Ebt13-5 | 7,209 | 412 | 6,616 | 311 |
| ***E. fitzii*** | Ef01-1 | 2,459 | 1,144 | 504 | 666 |
| | Ef01-2 | 36,284 | 3,208 | 27,977 | 2,992 |
| | Ef01-3 | 20,575 | 12,272 | 18,112 | 11,221 |
| | Ef01-4 | 21,007 | 3,040 | 19,382 | 2,819 |
| | Ef01-5 | 24,480 | 0 | 21,220 | 0 |
| ***E. lagascae*** | Ela07-1 | 46,652 | 7,949 | 38,755 | 7,498 |
| | Ela07-2 | 28,675 | 25,037 | 26,098 | 23,325 |
| | Ela07-3 | 43,375 | 44,502 | 43,522 | 40,688 |
| | Ela07-4 | 37,367 | 26,642 | 34,303 | 23,085 |
| | Ela07-5 | 21,787 | 68,778 | 18,544 | 49,696 |
| ***E. mediohispanicum*** | Em21-1 | 1,057 | 46 | 1,027 | 38 |
| | Em21-2 | 1,517 | 23 | 1,470 | 17 |
| | Em21-3 | 1,500 | 37 | 1,454 | 34 |
| | Em21-4 | 1,061 | 54 | 1,013 | 47 |
| | Em21-5 | 2,460 | 31 | 1,801 | 27 |

| | | | | | |
|---|---|---|---|---|---|
| | Em39-1 | 740 | 10 | 580 | 7 |
| | Em39-2 | 3,860 | 36 | 355 | 36 |
| | Em39-3 | 4,184 | 48 | 297 | 39 |
| | Em39-4 | 6,317 | 61 | 401 | 59 |
| | Em39-5 | 1,542 | 12 | 156 | 11 |
| | Em71-1 | 3,372 | 324 | 310 | 259 |
| | Em71-2 | 3,152 | 92 | 281 | 68 |
| | Em71-3 | 5,526 | 253 | 418 | 187 |
| | Em71-4 | 1,817 | 260 | 215 | 165 |
| | Em71-5 | 3,824 | 52 | 387 | 23 |
| ***E. nevadense*** | En05-1 | 19,550 | 3,660 | 22,540 | 3,260 |
| | En05-2 | 33,125 | 3,361 | 29,703 | 3,222 |
| | En05-3 | 20,691 | 38,519 | 19,623 | 36,417 |
| | En05-4 | 72,945 | 9,749 | 65,114 | 8,588 |
| | En05-5 | 101 | 6,507 | 72 | 5,839 |
| | En10-1 | 17,010 | 3,031 | 15,179 | 2,734 |
| | En10-2 | 15,650 | 70,323 | 13,255 | 65,241 |
| | En10-3 | 21,251 | 62,475 | 17,700 | 60,745 |
| | En10-4 | 32,198 | 66,797 | 30,531 | 63,176 |
| | En10-5 | 19,828 | 182,495 | 17,403 | 175,659 |
| | En12-1 | 14,249 | 101,910 | 13,363 | 100,098 |
| | En12-2 | 15,973 | 100,528 | 13,990 | 97,509 |
| | En12-3 | 18,005 | 6,829 | 16,465 | 6,298 |
| | En12-5 | 29,274 | 291,336 | 26,163 | 287,162 |
| ***E. popovii*** | Ep16-1 | 10,691 | 32,788 | 8,965 | 27,961 |
| | Ep16-2 | 2,753 | 26,133 | 2,590 | 15,418 |
| | Ep16-3 | 5,647 | 26,132 | 5,300 | 25,074 |
| | Ep16-4 | 6,081 | 18,519 | 4,967 | 17,651 |
| | Ep16-5 | 6,922 | 14,860 | 6,381 | 13,983 |
| | Ep20-1 | 19,099 | 55,464 | 13,841 | 48,638 |
| | Ep20-2 | 11,313 | 15,011 | 10,034 | 13,546 |
| | Ep20-3 | 4,570 | 7,092 | 4,108 | 5,309 |
| | Ep20-4 | 5,056 | 6,522 | 3,753 | 3,234 |
| | Ep20-5 | 5,861 | 25,346 | 5,119 | 22,381 |
| | Ep27-1 | 7,621 | 58,789 | 5,962 | 53,469 |
| | Ep27-2 | 6,239 | 44,639 | 5,227 | 36,287 |
| | Ep27-3 | 15,831 | 12,787 | 14,370 | 11,745 |
| | Ep27-4 | 6,207 | 38,248 | 5,147 | 35,449 |
| | Ep27-5 | 25,310 | 24,543 | 21,224 | 236 |

**Table S2.** Nucleotide and haplotype diversity for *E. baeticum* ITS1 and ITS2, at the three-level (species, population, individuals) analyzed.

| *E. baeticum* | Sample code | ITS1 | | ITS2 | |
|---|---|---|---|---|---|
| | | π | Hd | π | Hd |
| **Species level** | Ebb | 0.013 | 0.983 | 0.006 | 0.897 |
| **Population level** | Ebb07 | 0.015 | 1.000 | 0.009 | 0.933 |
| | Ebb10 | 0.012 | 0.963 | 0.004 | 1.000 |
| | Ebb12 | 0.015 | 1.000 | 0.008 | 0.933 |
| **Individual level** | Ebb07-1 | 0.022 | 1.000 | 0 | 0 |
| | Ebb07-2 | 0.011 | 1.000 | 0 | 0 |
| | Ebb07-3 | 0 | 0 | 0.010 | 1.00 |
| | Ebb07-4 | 0.012 | 1.000 | 0 | 0 |
| | Ebb07-5 | 0.018 | 1.000 | 0 | 0 |
| | Ebb10-1 | 0.018 | 1.000 | 0 | 0 |
| | Ebb10-2 | 0 | 0 | 0 | 0 |
| | Ebb10-3 | 0.011 | 1.000 | 0 | 0 |
| | Ebb10-4 | 0 | 0 | 0 | 0 |
| | Ebb10-5 | 0.017 | 1.000 | 0 | 0 |
| | Ebb12-1 | 0.009 | 1.000 | 0 | 0 |
| | Ebb12-2 | 0.011 | 1.000 | 0.008 | 1.000 |
| | Ebb12-3 | 0 | 0 | 0 | 0 |
| | Ebb12-4 | 0.015 | 1.000 | 0 | 0 |
| | Ebb12-5 | 0.011 | 1.000 | 0 | 0 |

**Table S3.** Nucleotide and haplotype diversity for *E. bastetanum,* ITS1 and ITS2 samples, at the three-level analyzed.

| *E. bastetanum* | Sample code | ITS1 | | ITS2 | |
|---|---|---|---|---|---|
| | | π | Hd | π | Hd |
| **Species level** | Ebt | 0.013 | 0.983 | 0.006 | 0.893 |
| **Population level** | Ebt01 | 0.013 | 0.969 | 0.005 | 0.694 |
| | Ebt12 | 0.012 | 0.991 | 0.008 | 0.900 |
| | Ebt13 | 0.013 | 0.975 | 0.005 | 0.916 |
| **Individual level** | Ebt01-1 | 0.013 | 1.000 | 0.005 | 0.694 |
| | Ebt01-2 | 0.014 | 1.000 | 0 | 0 |
| | Ebt01-3 | 0.020 | 1.000 | 0.008 | 1.000 |
| | Ebt01-4 | 0.019 | 1.000 | 0.021 | 1.000 |
| | Ebt01-5 | 0.017 | 1.000 | 0.005 | 0.666 |
| | Ebt12-1 | 0.015 | 1.000 | 0.010 | 1.000 |
| | Ebt12-2 | 0.015 | 1.000 | 0 | 0 |
| | Ebt12-3 | 0.015 | 1.000 | 0 | 0 |
| | Ebt12-4 | 0.013 | 1.000 | 0 | 0 |
| | Ebt12-5 | 0.011 | 1.000 | 0 | 0 |
| | Ebt13-1 | 0.013 | 1.000 | 0.004 | 1.000 |
| | Ebt13-2 | 0.018 | 1.000 | 0.004 | 1.000 |
| | Ebt13-3 | 0.017 | 1.000 | 0.006 | 1.000 |
| | Ebt13-4 | 0.013 | 1.000 | 0.005 | 1.000 |
| | Ebt13-5 | 0.030 | 1.000 | 0.006 | 1.000 |

**Table S4.** Nucleotide and haplotype diversity for *E. fitzii,* ITS1 and ITS2 samples, at the two level analyzed.

| ***E. fitzii*** | **Sample code** | **ITS1** | | **ITS2** | |
|---|---|---|---|---|---|
| | | **π** | **Hd** | **π** | **Hd** |
| **Species level** | Ef | 0.011 | 0.944 | 0.009 | 0.972 |
| **Individual level** | Ef01-1 | 0 | 0 | 0.012 | 1.000 |
| | Ef01-2 | 0 | 0 | 0.010 | 1.000 |
| | Ef01-3 | 0.008 | 1.000 | 0.008 | 1.000 |
| | Ef01-4 | 0 | 0 | 0.010 | 1.000 |
| | Ef01-5 | 0.019 | 1.000 | 0 | 0 |

**Table S5.** Nucleotide and haplotype diversity for *E. lagascae,* ITS1 and ITS2 samples, at the three-level analyzed.

| ***E. lagascae*** | **Sample code** | **ITS1** | | **ITS2** | |
|---|---|---|---|---|---|
| | | **π** | **Hd** | **π** | **Hd** |
| **Species level** | Ela | 0.011 | 1.00 | 0.005 | 0.733 |
| **Individual level** | Ela07-1 | 0 | 0 | 0 | 0 |
| | Ela07-2 | 0.008 | 1.00 | 0 | 0 |
| | Ela07-3 | 0 | 0 | 0 | 0 |
| | Ela07-4 | 0 | 0 | 0.007 | 1.000 |
| | Ela07-5 | 0.019 | 1.00 | 0 | 0 |

**Table S6.** Nucleotide and haplotype diversity for *E. mediohispanicum,* ITS1 and ITS2 samples, at the three-level analyzed.

| *E. mediohispanicum* | Sample code | ITS1 | | ITS2 | |
|---|---|---|---|---|---|
| | | π | Hd | π | Hd |
| **Species level** | Em | 0.300 | 0.969 | 0.003 | 0.805 |
| **Population level** | Em21 | 0.001 | 0.400 | 0.002 | 0.750 |
| | Em39 | 0.001 | 0.955 | 0.001 | 0.333 |
| | Em71 | 0.003 | 0.963 | 0.003 | 0.847 |
| **Individual level** | Em21-1 | 0.004 | 0.812 | 0.005 | 1.000 |
| | Em21-2 | 0 | 0 | 0.002 | 1.000 |
| | Em21-3 | 0 | 0 | 0.002 | 1.000 |
| | Em21-4 | 0 | 0 | 0 | 0 |
| | Em21-5 | 0 | 0 | 0 | 0 |
| | Em39-1 | 0.014 | 1.000 | 0.003 | 1.000 |
| | Em39-2 | 0 | 0 | 0 | 0 |
| | Em39-3 | 0 | 0 | 0 | 0 |
| | Em39-4 | 0.005 | 1.000 | 0 | 0 |
| | Em39-5 | 0.014 | 1.000 | 0 | 0 |
| | Em71-1 | 0.014 | 1.000 | 0.007 | 1.000 |
| | Em71-2 | 0.014 | 1.000 | 0.003 | 1.000 |
| | Em71-3 | 0 | 0 | 0.003 | 1.000 |
| | Em71-4 | 0.014 | 1.000 | 0.001 | 1.000 |
| | Em71-5 | 0.015 | 1.000 | 0.003 | 1.000 |

**Table S7.** Nucleotide and haplotype diversity for *E. nevadense,* ITS1 and ITS2 samples, at the three-level analyzed.

| ***E. nevadense*** | **Sample code** | **ITS1** | | **ITS2** | |
|---|---|---|---|---|---|
| | | **π** | **Hd** | **π** | **Hd** |
| **Species level** | En | 0.010 | 0.938 | 0.006 | 0.941 |
| **Population level** | En05 | 0.008 | 0.785 | 0.010 | 0.933 |
| | En10 | 0.004 | 0.800 | 0.007 | 1.000 |
| | En12 | 0.009 | 0.952 | 0.003 | 0.833 |
| **Individual level** | En05-1 | 0.008 | 1.000 | 0.007 | 1.000 |
| | En05-2 | 0 | 0 | 0 | 0 |
| | En05-3 | 0 | 0 | 0 | 0 |
| | En05-4 | 0.008 | 1.000 | 0 | 0 |
| | En05-5 | 0.011 | 1.000 | 0 | 0 |
| | En10-1 | 0 | 0 | 0 | 0 |
| | En10-2 | 0 | 0 | 0 | 0 |
| | En10-3 | 0 | 0 | 0 | 0 |
| | En10-4 | 0 | 0 | 0 | 0 |
| | En10-5 | 0.008 | 1.000 | 0 | 0 |
| | En12-1 | 0 | 0 | 0 | 0 |
| | En12-2 | 0.014 | 1.000 | 0 | 0 |
| | En12-3 | 0.011 | 1.000 | 0 | 0 |
| | En12-4 | 0 | 0 | 0 | 0 |
| | En12-5 | 0 | 0 | 0 | 0 |

**Table S8.** Nucleotide and haplotype diversity for E. popovii, ITS1 and ITS2 samples, at the three-level analyzed.

| *E. popovii* | Sample code | ITS1 | | ITS2 | |
|---|---|---|---|---|---|
| | | π | Hd | π | Hd |
| **Species level** | Ep | 0.015 | 0.984 | 0.007 | 0.943 |
| **Population level** | Ep16 | 0.013 | 0.977 | 0.007 | 0.977 |
| | Ep20 | 0.016 | 1.000 | 0.008 | 0.944 |
| | Ep27 | 0.004 | 0.888 | 0.004 | 0.866 |
| **Individual level** | Ep16-1 | 0 | 0 | 0.010 | 1.000 |
| | Ep16-2 | 0.014 | 1.000 | 0.011 | 1.000 |
| | Ep16-3 | 0.011 | 1.000 | 0 | 0 |
| | Ep16-4 | 0.019 | 1.000 | 0.005 | 1.000 |
| | Ep16-5 | 0.022 | 1.000 | 0.008 | 1.000 |
| | Ep20-1 | 0.019 | 1.000 | 0.010 | 1.000 |
| | Ep20-2 | 0.017 | 1.000 | 0.010 | 1.000 |
| | Ep20-3 | 0.014 | 1.000 | 0.005 | 1.000 |
| | Ep20-4 | 0.011 | 1.000 | 0.010 | 1.000 |
| | Ep20-5 | 0.018 | 1.000 | 0 | 0 |
| | Ep27-1 | 0 | 0 | 0 | 0 |
| | Ep27-2 | 0 | 0 | 0 | 0 |
| | Ep27-3 | 0.005 | 1.000 | 0 | 0 |
| | Ep27-4 | 0 | 0 | 0.008 | 1.000 |
| | Ep27-5 | 0.002 | 1.000 | 0 | 0 |

**Table S9.** Number of total haplotypes, frequency of each haplotype as relative abundance (based on the total of sequences after cd-hit analysis), number of haplotypes shared among different populations from the same species, and number of haplotypes shared among *E. baeticum* and the other *Erysimum* species studied here.

| ***E. baeticum*** | | ITS1 | | | | ITS2 | | | |
|---|---|---|---|---|---|---|---|---|---|
| | Sample | Number | Relative abundance (%) | Shared with other populations | Shared with other species | Number | Relative abundance (%) | Shared with other populations | Shared with other species |
| **Species level** | Ebb | 31 | H1: 10.25<br>H2: 7.69<br>H3-H5: 5.12<br>H6-H31: 2.56 | 3 | 0 | 9 | H1: 29.41<br>H2-H5: 11.76<br>H6-H9: 5.88 | 5 | 0 |
| **Population level** | Ebb07 | 13 | H1-H13: 7.69 | H1: Ebb12 | | 4 | H1-H2: 33.33<br>H3-H4: 16.66 | H1: Ebb12<br>H2: Ebb10, Ebb12 | |
| | Ebb10 | 9 | H1: 33.33<br>H2-H9: 8.33 | H1: Ebb12 | | 4 | H1: 40<br>H2-H4: 20 | H2: Ebb07, Ebb12<br>H3: Ebb12<br>H6: Ebb12 | |
| | Ebb12 | 11 | H1-H3: 14.28<br>H4-H11: 7.14 | H2: Ebb07<br>H3: Ebb10 | | 4 | H1: 50<br>H2-H4: 16.66 | H1: Ebb07<br>H2: Ebb12, Ebb07<br>H3: Ebb10<br>H4: Ebb10 | |
| **Individual level** | Ebb07-1 | 2 | H1: 64.55<br>H2: 11.84 | | | 1 | H1: 98.42 | | |
| | Ebb07-2 | 3 | H1: 51.38<br>H2: 24.58<br>H3: 12.89 | | | 2 | H1: 64.67<br>H2: 32.90 | | |
| | Ebb07-3 | 1 | H1: 90.12 | | | 1 | H1: 40.41 | | |
| | Ebb07-4 | 3 | H1: 31.89<br>H2: 27.79<br>H3: 12.08 | | | 1 | H1: 39.55 | | |
| | Ebb07-5 | 4 | H1: 54.89<br>H2: 16.00<br>H3: 6.18<br>H4: 6.02 | | | 1 | H1: 31.83 | | |
| | Ebb10-1 | 5 | H1: 31.01<br>H2: 18.22<br>H3: 10.23<br>H4: 8.29<br>H5: 6.82 | | | 1 | H1: 40.77 | | |
| | Ebb10-2 | 1 | H1: 93.17 | | | 1 | H1: 35.76 | | |
| | Ebb10-3 | 2 | H1: 80.78<br>H2: 10.93 | | | 1 | H1: 60.29 | | |
| | Ebb10-4 | 1 | H1: 91.02 | | | 1 | H1: 43.22 | | |
| | Ebb10-5 | 3 | H1: 61.98<br>H2: 17.56<br>H3: 5.18 | | | 1 | H1: 40.02 | | |
| | Ebb12-1 | 3 | H1: 55.44<br>H2: 19.68<br>H3: 12.30 | | | 2 | H1: 22.50<br>H2: 15.69 | | |
| | Ebb12-2 | 3 | H1: 61.30<br>H2: 11.94<br>H3: 8.87 | | | 1 | H1: 43.72 | | |
| | Ebb12-3 | 1 | H1: 85.40 | | | 1 | H1: 37.98 | | |
| | Ebb12-4 | 5 | H1: 38.78<br>H2: 15.61<br>H3: 15.20<br>H4: 12.68<br>H5: 7.56 | | | 1 | H1: 40.88 | | |
| | Ebb12-5 | 2 | H1: 42.54<br>H2: 38.41 | | | 1 | H1: 41.50 | | |

**Table S10.** Number of total haplotypes, frequency of each haplotype (based on the total of sequences after cd-hit analysis), number of haplotypes shared among different populations from the same species, and number of haplotypes shared among *E. bastetanum* and other *Erysimum* species studied here.

| ***E. bastetanum*** | | **ITS1** | | | | **ITS2** | | | |
|---|---|---|---|---|---|---|---|---|---|
| | **Sample** | **Number** | **Relative abundance (%)** | **Shared with other populations** | **Shared with other species** | **Number** | **Relative abundance (%)** | **Shared with other populations** | **Shared with other species** |
| **Species level** | Ebt | 38 | H1: 10.86<br>H2: 6.52<br>H3: 4.34<br>H4-H38: 2.17 | 2 | 0 | 17 | H1: 27.5<br>H2: 15<br>H3-H4: 10<br>H5-H6: 5<br>H7-H17: 2.5 | 2 | 0 |
| **Population level** | Ebt01 | 9 | H1: 25<br>H2: 16.6<br>H3-H9: 8.33 | H1:Ebt12, Ebt13<br>H2: Ebt13 | | 4 | H1: 55.55<br>H2: 22.22<br>H3-H4: 11.11 | H1: Ebt12 | |
| | Ebt12 | 15 | H1: 12.5<br>H2: 6.25<br>H3-H15: 6.25 | H1: Ebt01, Ebt13 | | 4 | H1: 40<br>H2-H4: 20 | H1: Ebt01<br>H2: Ebt13 | |
| | Ebt13 | 13 | H1: 5.55<br>H2-H13: 5.55 | H1: Ebt01, Ebt12<br>H2: Ebt01 | | 4 | H1: 60.25<br>H2: 17.32<br>H3: 15.94<br>H4: 3.82 | H2: Ebt12 | |
| **Individual level** | Ebt01-1 | 3 | H1: 43.75<br>H2: 28.68<br>H3: 9.5 | | | 1 | H1: 95.36 | | |
| | Ebt01-2 | 2 | H1: 59.43<br>H2: 23.67 | | | 1 | H1: 95.61 | | |
| | Ebt01-3 | 2 | H1: 61.91<br>H2: 29.94 | | | 2 | H1: 83.54<br>H2: 14.36 | | |
| | Ebt01-4 | 2 | H1: 74.73<br>H2: 20.69 | | | 2 | H1: 91.87<br>H2: 5.51 | | |
| | Ebt01-5 | 3 | H1: 46.94<br>H2: 22.18<br>H3: 10.46 | | | 2 | H1: 86.81<br>H2: 11.74 | | |
| | Ebt12-1 | 3 | H1: 75.14<br>H2: 9.05<br>H3: 5.91 | | | 3 | H1: 59.53<br>H2: 31.10<br>H3: 7.31 | | |
| | Ebt12-2 | 3 | H1: 41.94<br>H2: 21.94<br>H3: 11.91 | | | 1 | H1: 91.82 | | |
| | Ebt12-3 | 4 | H1: 33.87<br>H2: 17.93<br>H3: 11.81<br>H4: 10.43 | | | 1 | H1: 96.04 | | |
| | Ebt12-4 | 4 | H1: 33.38<br>H2: 25.34<br>H3: 15.49<br>H4: 5 | | | 0 | 0 | | |
| | Ebt12-5 | 2 | H1: 84.18<br>H2: 8.45 | | | 0 | 0 | | |
| | Ebt13-1 | 3 | H1: 55.78<br>H2: 23.92<br>H3: 9.36 | | | 5 | H1: 30<br>H2: 27.27<br>H3: 21.21<br>H4: 15.15<br>H5: 6 | | |
| | Ebt13-2 | 5 | H1: 38.84<br>H2: 20.40<br>H3: 9.66<br>H4: 7.66<br>H5: 6.16 | | | 4 | H1: 46<br>H2: 22.10<br>H3: 18.94<br>H4: 12.63 | | |
| | Ebt13-3 | 2 | H1: 53.444<br>H2: 36.14 | | | 3 | H1: 57.02<br>H2: 26.40<br>H3: 16.57 | | |
| | Ebt13-4 | 3 | H1: 34.01<br>H2: 27.41<br>H3: 25.92 | | | 3 | H1: 63.33<br>H2: 24.44<br>H3: 12.22 | | |
| | Ebt13-5 | 5 | H1: 37.57<br>H2: 31.77 | | | 3 | H1: 72.34<br>H2: 14.14 | | |

| | |
|---|---|
| H3: 10.75<br>H4: 5.85<br>H5: 5.81 | H3: 9.6 |

**Table S11.** Number of total haplotypes, frequency of each haplotype (based on the total of sequences after cd-hit analysis), and number of haplotypes shared among *E. fitzii* and other *Erysimum* species studied here.

| ***E. fitzii*** | | **ITS1** | | | **ITS2** | | |
|---|---|---|---|---|---|---|---|
| | **Sample** | **Number** | **Relative abundance (%)** | **Shared with other species** | **Number** | **Relative abundance (%)** | **Shared with other species** |
| **Species level** | Ef01 | 6 | H1-H3: 22.22<br>H4-H6: 11.11 | 0 | 7 | H1-H7: 14.27 | 0 |
| **Individual level** | Ef01-1 | 1 | H1: 20 | | 1 | H1: 83.07 | |
| | Ef01-2 | 4 | H1: 27.42<br>H2: 24.04<br>H3: 20.34<br>H4: 5.27 | | 1 | H1: 91.01 | |
| | Ef01-3 | 1 | H1: 88.02 | | 2 | H1: 51.63<br>H2: 42.97 | |
| | Ef01-4 | 1 | H1: 92.26 | | 2 | H1: 81.30<br>H2: 10.50 | |
| | Ef01-5 | 2 | H1: 58.84<br>H2: 27.83 | | 1 | H1: 85.16 | |

**Table S12.** Number of total haplotypes, frequency of each haplotype (based on the total of sequences after cd-hit analysis), and number of haplotypes shared among *E. lagascae* and other *Erysimum* species studied here.

| *E. lagascae* | | ITS1 | | | ITS2 | | |
|---|---|---|---|---|---|---|---|
| | Sample | Number | Relative abundance (%) | Shared with other species | Number | Relative abundance (%) | Shared with other species |
| **Species level** | Ela07 | 7 | H1-H7: 14.27 | 0 | 2 | H1: 74.98<br>H2: 19.34 | 0 |
| **Individual level** | Ela07-1 | 1 | H1: 83.07 | | 1 | H1: 93.16 | |
| | Ela07-2 | 1 | H1: 91.01 | | 1 | H1: 91.42 | |
| | Ela07-3 | 2 | H1: 57.63<br>H2: 42.97 | | 1 | H1: 97.55 | |
| | Ela07-4 | 2 | H1: 81.30<br>H2: 10.50 | | 2 | H1: 46.94<br>H2: 25.31 | |
| | Ela07-5 | 1 | H1: 85.16 | | 1 | H1: 94.5 | |

**Table S13.** Number of total haplotypes, frequency of each haplotype as relative abundance (based on the total of sequences after cd-hit analysis), number of haplotypes shared among different populations from the same species, and number of haplotypes shared among *E. mediohispanicum* and the other *Erysimum* species studied here.

| ***E. mediohispanicum*** | | **ITS1** | | | | **ITS2** | | | |
|---|---|---|---|---|---|---|---|---|---|
| | **Sample** | **Number** | **Relative abundance (%)** | **Shared with other populations** | **Shared with other species** | **Number** | **Relative abundance (%)** | **Shared with other populations** | **Shared with other species** |
| **Species level** | Em | 10 | H1: 23.67<br>H2: 22.79<br>H3: 20.25<br>H4: 12.54<br>H5: 9.92<br>H6:2.92<br>H7: 2.41<br>H8-H10: 1.5 | 1 | 5 | 13 | H1: 37.70<br>H2: 18.80<br>H3: 16.07<br>H4: 10.19<br>H5: 4<br>H6: 4.14<br>H7: 1.98<br>H8: 1.73<br>H9-H13: 1 | 0 | 0 |
| **Population level** | Em21 | 2 | H1: 63.79<br>H2: 36.20 | | | 4 | H1: 44.85<br>H2: 36.02<br>H3: 11.74<br>H4: 5.14 | | |
| | Em39 | 3 | H1: 19<br>H2: 14<br>H3: 14 | H1: Em71 | H1: Ebt12, Ebt13, Ebt01, En12<br>H5: En10 | 1 | H1: 96.64 | | |
| | Em71 | 5 | H1-H3: 10<br>H4-H5: 7 | H1: Em39 | H1: Ebt12, Ebt13, Ebt01, En12<br>H2: Ebt01, Ebt12, Ebt13<br>H3: Ebt13<br>H4: Ebt13 | 5 | H1: 25.92<br>H2: 19.65<br>H3: 18.66<br>H4: 17.52<br>H5: 7.12 | | |
| **Individual level** | Em21-1 | 1 | H1: 97.16 | | | 4 | H1: 39.47<br>H2: 31.57<br>H3: 21.05<br>H4: 7.89 | | |
| | Em21-2 | 1 | H1: 96.90 | | | 2 | H1: 70.58<br>H2: 29.41 | | |
| | Em21-3 | 1 | H1: 96.93 | | | 3 | H1: 58.82<br>H2: 32.35<br>H3: 8.82 | | |
| | Em21-4 | 2 | H1: 65.97<br>H2: 29.50 | | | 4 | H1: 63.82<br>H2: 19.14<br>H3: 8.51<br>H4: 8.51 | | |
| | Em21-5 | 1 | H1: 73.21 | | | 1 | H1: 100 | | |
| | Em39-1 | 3 | H1: 51.03<br>H2: 39.31<br>H3: 9.65 | | | 2 | H1: 55.55<br>H2: 44.44 | | |
| | Em39-2 | 5 | H1: 33.34<br>H2: 28.14<br>H3: 17.51<br>H4: 12.45<br>H5: 8.51 | | | 2 | H1: 57.14<br>H2: 42.85 | | |
| | Em39-3 | 2 | H1: 85.52<br>H2: 14.47 | | | 8 | H1: 80.55<br>H2-H8: 2.77 | | |
| | Em39-4 | 1 | H1: 100 | | | 1 | H1: 100 | | |
| | Em39-5 | 4 | H1: 48.44<br>H2: 25.77<br>H3: 16.14<br>H4: 9.62 | | | 1 | H1: 100 | | |
| | Em71-1 | 4 | H1: 52.22<br>H2: 36.75<br>H3: 5.55<br>H4: 5.54 | | | 1 | H1: 100 | | |
| | Em71-2 | 4 | H1: 48.06<br>H2: 24.11<br>H3: 18.49<br>H4: 9.32 | | | 4 | H1: 38.61<br>H2: 31.27<br>H3: 18.91<br>H4: 11.19 | | |
| | Em71-3 | 3 | H1: 46.24<br>H2: 39.33 | | | 3 | H1: 52.94<br>H2: 29.41 | | |

| | | | | |
|---|---|---|---|---|
| | | H3: 13.77 | | H3: 17.64 |
| Em71-4 | 2 | H1: 74 | 5 | H1: 25.66 |
| | | H2: 26 | | H2: 23.52 |
| | | | | H3: 20.32 |
| | | | | H4: 17.64 |
| | | | | H5: 12.83 |
| Em71-5 | 3 | H1: 44.27 | 4 | H1: 49.69 |
| | | H2: 41.42 | | H2: 23.63 |
| | | H3: 14.30 | | H3: 13.93 |
| | | | | H4: 12.72 |

**Table S14.** Number of total haplotypes, frequency of each haplotype as relative abundance (based on the total of sequences after cd-hit analysis), number of haplotypes shared among different populations from the same species, and number of haplotypes shared among *E. nevadense* and the other *Erysimum* species studied here.

| ***E. nevadense*** | | ITS1 | | | | ITS2 | | | |
|---|---|---|---|---|---|---|---|---|---|
| | Sample | Number | Relative abundance (%) | Shared with other populations | Shared with other species | Number | Relative abundance (%) | Shared with other populations | Shared with other species |
| **Species level** | En | 15 | H1: 23,80<br>H2: 14.28<br>H3-H15: 4.7 | 4 | H1: Ebt. Em<br>H47: Ebt<br>H66: Ef<br>H81: Em | 12 | H1-H2: 16.66<br>H3-H4: 11.11<br>H5-H12: 5.55 | 4 | 0 |
| **Population level** | En05 | 5 | H1: 50<br>H2: 12.5<br>H3: 12.5<br>H4: 12.5<br>H5: 12.5 | H1,H2: En10<br>H3: En12 | H1: Ebt12<br>H2: Ef<br>H4: En12, Em71, Em39, Ebt13, Ebt12, Ebt01 | 6 | H1: 28.57<br>H2-H6: 14.28 | H1: En10<br>H2: En12<br>H4: En12 | |
| | En10 | 4 | H1: 50<br>H2: 16.66<br>H3: 16.66<br>H4: 16.66 | H1,H2: En05<br>H4: En12 | H3: Em39 | 7 | H1-H7: 14.28 | H1: En05<br>H3: En12 | |
| | En12 | 7 | H1-H7: 14.28 | H4: En10<br>H3: En05 | H4: En05. Em71. Em39. Ebt13. Ebt12. Ebt01 | 3 | H1: 50<br>H2-H3: 25 | H2: En05<br>H4: En05 | |
| **Individual level** | En05-1 | 1 | H1: 86.73 | | | 2 | H1: 77.20<br>H2: 11.88 | | |
| | En05-2 | 2 | H1: 76.73<br>H2: 12.93 | | | 1 | H1: 91.81 | | |
| | En05-3 | 1 | H1: 93.09 | | | 1 | H1: 94.54 | | |
| | En05-4 | 2 | H1: 59.24<br>H2: 30.02 | | | 1 | H1: 95.78 | | |
| | En05-5 | 2 | H1: 58.41<br>H2: 12.87 | | | 1 | H1: 89.73 | | |
| | En10-1 | 1 | H1: 89.23 | | | 2 | H1: 78.78<br>H2: 11.41 | | |
| | En10-2 | 1 | H1: 84.69 | | | 1 | H1: 92.77 | | |
| | En10-3 | 1 | H1: 83.29 | | | 1 | H1: 97.23 | | |
| | En10-4 | 1 | H1: 94.82 | | | 1 | H1: 94.57 | | |
| | En10-5 | 2 | H1: 53.33<br>H2: 34.43 | | | 2 | H1: 61.68<br>H2: 34.57 | | |
| | En12-1 | 1 | H1: 93.78 | | | 1 | H1: 98.22 | | |
| | En12-2 | 2 | H1: 76.02<br>H2: 11.55 | | | 1 | H1: 96.99 | | |
| | En12-3 | 3 | H1: 39.48<br>H2: 26.18<br>H3: 25.77 | | | 1 | H1: 92.22 | | |
| | En12-4 | - | | | | | | | |
| | En12-5 | 1 | H1: 89.37 | | | 1 | H1: 98.56 | | |

**Table S15.** Number of total haplotypes, frequency of each haplotype (based on the total of sequences after cd-hit analysis), number of haplotypes shared among different populations from the same species, and number of haplotypes shared among *E. popovii* and other *Erysimum* species studied here.

| ***E. popovii*** | | ITS1 | | | | ITS2 | | | |
|---|---|---|---|---|---|---|---|---|---|
| | Sample | Number | Relative abundance (%) | Shared with other populations | Shared with other species | Number | Relative abundance (%) | Shared with other populations | Shared with other species |
| **Species level** | Ep | 30 | H1: 8.5<br>H2: 8.5<br>H3: 5.7<br>H4-H30: 2.85 | 1 | 0 | 19 | H1: 18.51<br>H2: 14.81<br>H3-H19: 3.70 | 1 | 0 |
| **Population level** | Ep16 | 12 | H1-H12: 8.33 | | | 9 | H1: 20<br>H2-H9: 10 | H1: Ep20, Ep27 | |
| | Ep20 | 12 | H1: 15.38<br>H2-H12: 7.69 | H1: Ep27 | | 9 | H1-H2: 18.18<br>H3-H9: 9 | H1: Ep27, Ep16 | |
| | Ep27 | 7 | H1: 30<br>H2: 20<br>H3-H7: 10 | H1: Ep20 | | 4 | H1-H2: 33.33<br>H3-H4: 16.66 | H1: Ep16, Ep20 | |
| **Individual level** | Ep16-1 | 2 | H1: 65.55<br>H2: 18.30 | | | 4 | H1: 42.33<br>H2: 18.43<br>H3: 16.87<br>H4: 7.64 | | |
| | Ep16-2 | 2 | H1: 61.09<br>H2: 32.98 | | | 3 | H1: 25.23<br>H2: 21.94<br>H3: 11.82 | | |
| | Ep16-3 | 2 | H1: 62.93<br>H2: 30.91 | | | 1 | H1: 95.95 | | |
| | Ep16-4 | 2 | H1: 75.46<br>H2: 6.21 | | | 2 | H1: 85.98<br>H2: 9.33 | | |
| | Ep16-5 | 4 | H1: 32.72<br>H2: 27.70<br>H3: 26.19<br>H4: 5.56 | | | 2 | H1: 87.94<br>H2: 6.15 | | |
| | Ep20-1 | 2 | H1: 44.87<br>H2: 27.59 | | | 2 | H1: 78.89<br>H2: 8.79 | | |
| | Ep20-2 | 3 | H1: 72.67<br>H2: 8.49<br>H3: 7.52 | | | 2 | H1: 78.82<br>H2: 11.41 | | |
| | Ep20-3 | 2 | H1: 80.70<br>H2: 9.19 | | | 2 | H1: 67.72<br>H2: 7.13 | | |
| | Ep20-4 | 3 | H1: 48.23<br>H2: 20.90<br>H3: 5.08 | | | 2 | H1: 61.34<br>H2: 28.57 | | |
| | Ep20-5 | 3 | H1: 61.28<br>H2: 14.43<br>H3: 11.61 | | | 1 | H1: 88.30 | | |
| | Ep27-1 | 1 | H1: 78.23 | | | 1 | H1: 90.95 | | |
| | Ep27-2 | 2 | H1: 78.36<br>H2: 5.41 | | | 2 | H1: 41.02<br>H2: 40.26 | | |
| | Ep27-3 | 3 | H1: 49.91<br>H2: 27.51<br>H3: 13.34 | | | 2 | H1: 67.41<br>H2: 24.43 | | |
| | Ep27-4 | 2 | H1: 75.88<br>H2: 7.04 | | | 2 | H1: 84.08<br>H2: 8.59 | | |
| | Ep27-5 | 2 | H1: 69.96<br>H2: 13.89 | | | 2 | H1: 90.20<br>H2: 6.19 | | |
